# Can pharmacological enhancement of the placebo effect be a novel therapy for working memory impairments?

**DOI:** 10.1101/416123

**Authors:** Weihua Zhao, Benjamin Becker, Shuxia Yao, Xiaole Ma, Juan Kou, Keith M Kendrick

## Abstract

Working memory is considered as a core aspect of cognitive function and its impairment in a wide range of mental disorders has resulted in it being considered as an important transdiagnostic feature. To date pharmacological and behavioural strategies for augmenting working memory have achieved only moderate success. Here we have taken a different approach by combining expectancy effects with intranasal oxytocin as an adjunct given previous evidence that it may enhance placebo effects. In a randomised controlled clinical trial we demonstrate that while working memory performance is not influenced by expectancy per se when it is given in conjunction with oxytocin performance in terms of accuracy can be significantly enhanced following positive expectancy induction (placebo effect) and impaired following negative expectancy induction (nocebo effect). Thus combining expectancy effects with intranasal oxytocin may represent a radical new approach for improving working memory function in mental disorders.

## Introduction

Working memory represents a cognitive function that allows information to be held temporarily in mind and used operationally (Baddeley & Hitch, 1994). This core cognitive domain represents a critical building block for all higher order cognitive functions, including language, social interaction and decision making that critically guide our daily life. Working memory capacity is limited and in accordance with the critical contribution of this function to processes that guide our everyday life, working memory capacity is highly predictive of academic success and socio-economic status but also more general aspects of quality of life (Diamond, 2013).

Impairments in the domain of working memory have been observed across mental) disorders and strongly predict treatment success (Goodkind et al., 2015; Millan et al., 2012). Deficits in this cardinal cognitive domain are not targeted by traditional treatments and not only impede functional recovery but also critically preclude the efficacy of psychotherapeutic interventions by interfering with the adaptive learning process (Millan et al., 2012; Normann et al., 2012). The pharmacological enhancement of working memory to augment the efficacy of learning-based interventions has therefore received considerable attention during recent years (e.g. Normann et al., 2012).

Major efforts have been made from both, academic science as well as industry to develop novel approaches to enhance working memory capacity in order to boost general cognitive performance in the healthy individuals and treatment success in patients. However, despite concerted efforts and massive investments from pharmacological and cognitive training companies attempts at improving working memory capacity have proven to be a real challenge and results so far have been rather sobering. Traditional pharmacological and training-based approaches have been shown to produce only moderate improvements, often limited to individuals with low baseline performance or sleep deprivation, or no benefits at all when tested in controlled experiments (Repantis et al., 2010; Dresler et al., 2013; Kable et al., 2017).

Placebo effects in the form of creating psychological expectancies strongly modulate therapeutic outcome and can have positive effects in some domains, but studies to date have failed to demonstrate expectancy-induced cognitive enhancement per se (Schwarz et al., 2015). Recent findings that the neuropeptide oxytocin can enhance placebo-induced non-cognitive effects, and acceptance of expert advice (Kessner et al., 2013; Colloca et al., 2016; Luo et al., 2017), suggest its potential use as an adjunct to expectancy-induction in the cognitive domain. Against this background the present study aimed at evaluating whether intranasal oxytocin in combination with expectancy-induction has the potential to improve working memory.

## Methods

### Participants

To this end we conducted a randomized, between-subject placebo-controlled intranasal oxytocin (24IU) proof-of-concept study with four experimental arms (total n = 224, healthy males). All subjects were free from a current or history of neurological or psychiatric disorders as well as regular or current use of nicotine, alcohol, or other psychoactive substances. Written informed consent was obtained from all participants. Prior to the experiment all subjects completed validated questionnaires to control for confounding effects of depression (Beck Depression Inventory II, BDI II, Beck et al., 1996); anxiety (State-Trait Anxiety Inventory, STAI, Spielberger et al., 1983) and interpersonal trust (Interpersonal Trust Scale, ITS, Rotter, 1967).

The study and experimental protocols had full approval by the local ethics committee of the University of Electronic Science and Technology of China and adhered to the latest revision of the Declaration of Helsinki. Protocols and primary outcomes were pre-registered at clinicaltrials.gov (ID, NCT02745522).

### Expectancy induction

In line with previous research on the pharmacological enhancement of expectancy effects, standardized verbal instructions were employed for expectancy-induction (Colloca et al., 2016). Subjects were given different instructions as follows across four independent experimental arms: they received either no expectancy-induction (neutral information, Experiment 1), placebo (‘oxytocin enhances performance’, Experiment 2) or nocebo effect induction (‘oxytocin impairs performance’, Experiment 3) which served as an active comparator arm. Verbal instructions were given according to a standardized protocol employed by the same female experimenter dressed in a white coat and blinded for the treatment (oxytocin versus placebo) the subjects received. Given that oxytocin’s facilitation of acceptance of advice can be influenced by the gender of the expert (Luo et al., 2017), Experiment 2 was repeated with a male experimenter (Experiment 4) (see also Table 1).

**Table 1.** Experimental design and primary outcome measures

|  | Experiment 1 (n = 66) |  | Experiment 2 (n = 53) |  | Experiment 3 (n =53) |  |
| --- | --- | --- | --- | --- | --- | --- |
| Design |  |  |  |  |  |  |
| Expectancy-induction | Neutral (expectancy: none) |  | Placebo (expectancy: improve) |  | Nocebo (expectancy: impair) |  |
| Experimenter | Female (white coat) |  | Female (white coat) |  | Female (white coat) |  |
| Treatment | Placebo | Oxytocin | Placebo | Oxytocin | Placebo | Oxytocin |
| Groups | (n = 34) | (n = 32) | (n = 26) | (n = 27) | (n = 27) | (n = 26) |
| Results (Primary outcome - % difference in N-back task performance between oxytocin and placebo groups<sup>a</sup>) |  |  |  |  |  |  |
|  | Mean (SEM) | *p* | Mean (SEM) | *p* / *E.S.* | Mean (SEM) | *p* / *E.S.* |
| Verbal WM | .20 (1.3) | .86 | 5.3 (1.4) | .001 / .93 | -4.1 (1.4) | .005 / .97 |
| Spatial WM | .80 (1.4) | .60 | 3.6 (1.6) | .02 / .60 | -3.8 (1.6) | .02 / .63 |
| Social WM | -2.5 (1.9) | .18 | 4.6 (2.1) | .03 / .82 | -5.0 (2.1) | .02 / .65 |

Abbreviations: WM, working memory; E.S., Effect size (Cohen's)
<sup>a</sup> averaged accuracy difference across working memory load conditions (1-back and 2- back) reported given that no load-dependent interaction effects were found.

### Treatment

Subjects were randomly allocated to receive either a single intranasal dose of 24 IU OXT (3 puffs of 4IU per nostril with 30s between each puff – Sichuan Meike, Pharmaceutical Co., China) or placebo (PLC) with identical ingredients (also 3 puffs per nostril) other than the peptide. Treatment was administered in line with a standardized protocol (Guastella et al., 2013). Working memory assessment started 45 minutes after treatment administration.

### Primary outcome

Working memory accuracy, defined as percent correct responses, served as primary outcome. Working memory performance was assessed using a visual n-back task (Owen et al., 2005), a classical working memory paradigm during which subjects are presented a sequence of stimuli and have to indicate whether the current stimulus corresponds to the one that was shown ‘n’ trials before. To promote a comprehensive characterization of the treatment effects, performance was assessed in the domains of spatial, verbal and social working memory and at two levels of working memory load (1-back, 2-back) (details see Owen et al., 2005; Becker et al., 2010; Neta & Whalen, 2011). Domain- and load-specific performance was acquired in separate blocks presented in a counterbalanced order. Each block was preceded by a cue indicating the working memory domain and working memory load of the block.

## Results

Examination of potential confounders revealed that the intervention groups did not differ with respect to important confounders including age, education, depression, anxiety and trust (all *ps* > 0.12). Initial analysis of the primary outcome did not reveal load-dependent effects of expectancy-induction and treatment on accuracy (all *ps* > 0.22), consequently the factor load was discarded from subsequent analyses. Evaluating the effects of oxytocin on expectancy-induced modulation of working memory using domain-specific ANOVAs, with expectancy-induction (neutral, enhancement, impairment) and treatment (oxytocin, placebo) as between-subject factors (Experiments 1-3), revealed that neither treatment nor expectancy-induction per se influenced performance (all *ps* > 0.11). Importantly, significant expectancy-induction x treatment interactions were consistently observed across all working memory domains (verbal, *F*(_2, 166_) = 10.70, *p* < 0.001, *η*^2^_p_ = 0.11; spatial, *F*(_2, 166_) = 5.52, *p* = 0.005, *η*^2^_p_ = 0.06; social, *F*(_2, 166_) =5.79, *p* = 0.004, *η*^2^_p_ = 0.07). Post-hoc, Bonferroni-corrected tests indicated that oxytocin produced both expectancy-induced enhancement (verbal, +5%, 95%-CI, 3 to 8%; spatial, +4%,95%-CI, 1 to 7%; social, +5%, 95%-CI, 1 to 9%) and impairment (verbal -4%, 95%-CI, -7 to -1%; spatial, -4%, 95%-CI, -7 to -1%; social, -5%, 95%-CI, -9 to -1%) of working memory with medium or large effect sizes (effect size > 0.60 in all cases, details see Table 1, Figure 1). Next, the potential role of experimenter gender was investigated using domain-specific ANOVAs including treatment and gender as between-subject factors (Experiment 2, 4). This revealed that oxytocin produced an expectancy-induced working memory enhancement independent of experimenter gender (treatment main effect, verbal, *F*_(1, 101)_ = 16.29, *p* < 0.001, *η*^2^_p_ = 0.14; spatial, *F*_(1, 101)_ = 12.08, *p* = 0.001, *η*^2^_p_ = 0.11; social, *F*_(1, 101)_ = 10.81, *p* = 0.001, *η*^2^_p_ = 0.10; all treatment x gender interactions *ps* > 0.21).

**Figure 1.**
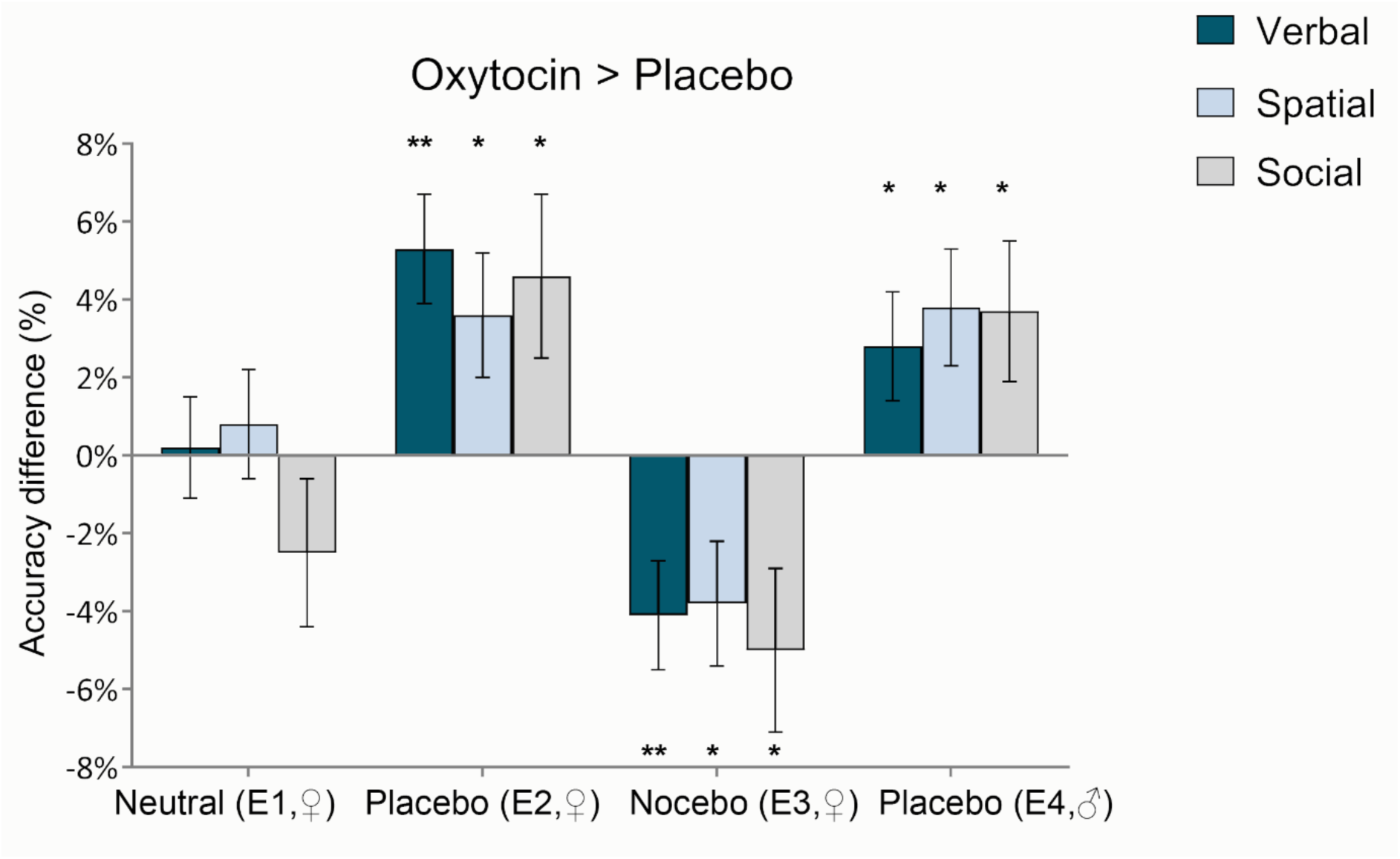
Oxytocin-induced expectancy-driven modulation of working memory ^a^. ^a^ Differences between the oxytocin and the placebo group in the four experiments (neutral, placebo effect, nocebo effect and placebo effect by a male experimenter) in terms of verbal, spatial and social n-back tasks are displayed (oxytocin > placebo). * *p* <.05; ** *p* <.01,Bonferroni-corrected post hoc comparison; ♀ female experimenter; ♂ male experimenter; E1-4, experimental arms 1-4 conducted in independent samples.

## Discussion

In summary, the present findings demonstrate for the first time that adjunct oxytocin treatment induces expectancy-driven enhancement of working memory. In line with previous findings neither expectancy-induction nor oxytocin per se affected working memory performance (Schwarz et al., 2015). Together with the observation that effects were concordant with the direction of expectancy induction (i.e. improvement vs. impairment) this points to oxytocinergic enhancement of expectancy-induction as an underlying mechanism that drives the modulation of working memory. With respect to previous pharmacological and training-based strategies for cognitive enhancement it is noteworthy that the effects were observed in a sample with high baseline performance and without sleep deprivation (Repantis et al., 2010; Dresler et al., 2010).

Although the present study did not acquire additional neurobiological indices it is important to consider the possible underlying neural mechanisms. Whereas previous research indicates that the endogenous opioid system strongly contributes to placebo analgesia, studies that examined expectancy-induced improvement of motor performance and positive drug effects suggest that effects in these domains are mediated by the dopaminergic system and modulation of striato-frontal circuits (Bendetti & Amanzio, 2013; Kaasinen et al., 2004; Volkow et al., 2006). Dopaminergic neurotransmission in these pathways has not only been associated with the strengths of the placebo response (Scott et al., 2007), but also critically differentiated between the placebo and nocebo responses (Scott et al., 2008) and contributes to working memory performance and cognitive effort (Diamond, 2007, Westbrook & Braver, 2016). Recent animal models suggest that oxytocin promotes social interactions via direct effects on dopaminergic neurotransmission in these pathways (Hung et al., 2017), which together with previous findings on an oxytocinergic amplification of expectancy-related processing in the underlying pathways in humans (Scheele et al., 2014; Kreuder et al., 2017) may suggest a contribution of oxytocin-dopamine interactions on the present findings.

Translated into the clinical context, augmentation strategies with adjunct oxytocin treatment may help to boost therapeutic efficacy and functional recovery. To this end, combining oxytocin and expectancy-induction could provide a radical innovative strategy to overcome working memory deficits in patients with mental disorders.

## Acknowledgments

This work was supported by grants from National Natural Science Foundation of China (NSFC) [31530032; 91632117].

